# Hippocampal Transplants of Fetal GABAergic Progenitors Regulate Adult Neurogenesis in Mice with Temporal Lobe Epilepsy

**DOI:** 10.1101/2022.01.11.475674

**Authors:** Muhammad N. Arshad, Simon Oppenheimer, Jaye Jeong, Bilge Buyukdemirtas, Janice R. Naegele

## Abstract

GABAergic interneurons within the dentate gyrus of the hippocampus regulate adult neurogenesis, including proliferation, migration, and maturation of new granule cells born in the subgranular zone (SGZ) of the dentate gyrus (DG). In temporal lobe epilepsy (TLE), some adult-born granule cells migrate ectopically into the hilus, and these cells contribute to increased hyperexcitability and seizures. Yet, transplanting embryonic day 13.5 fetal mouse medial ganglionic eminence (MGE) GABAergic progenitors into the hippocampus of mice with TLE ameliorates spontaneous seizures, due in part, to increased postsynaptic inhibition of adult-born granule cells. Here, we asked whether MGE progenitor transplantation affects earlier stages of adult neurogenesis, by comparing patterns of neurogenesis in naïve mice and epileptic (TLE) mice, with or without MGE transplants. In naïve and TLE mice, transplanted MGE cells showed comparable migration and process outgrowth. However, in TLE mice with MGE transplants, fewer adult-born Type 3 progenitors migrated ectopically. Furthermore, more Type 3 progenitors survived and migrated into the granule cell layer (GCL), as determined by immunostaining for doublecortin or the thymidine analogue, bromodeoxyuridine (BrdU). To determine whether MGE transplants affected earlier stages of adult neurogenesis, we compared proliferation in the SGZ two-hours after pulse labeling with BrdU in Naive vs. TLE mice and found no significant differences. Furthermore, MGE progenitor transplantation had no effect on cell proliferation in the SGZ. Moreover, when compared to naïve mice, TLE mice showed increases in inverted Type 1 progenitors and Type 2 progenitors, concomitant with a decrease in the normally oriented radial Type 1 progenitors. Strikingly, these alterations were abrogated by MGE transplantation. Thus, MGE transplants appear to reverse seizure-induced abnormalities in adult neurogenesis by increasing differentiation and radial migration of adult-born granule cell progenitors, outcomes that may ameliorate seizures.

## INTRODUCTION

Adult dentate neurogenesis is a dynamic process affected by age, physical exercise, hormones, chronic stress, and neurological disorders (Eisch and Petrik, 2012; Lucassen et al., 2015; Vivar et al., 2016; Kuhn et al., 2018; Voss et al., 2019; Gage, 2021). In the rodent brain, the process occurs in the neurogenic niche of the dentate gyrus, called the subgranular zone (SGZ). This thin layer of cells between the granule cell layer (GCL) and the hilus includes a neural stem cell pool and provides an environment suitable for neural stem cell proliferation. Type 1 or radial glia-like (RGL) progenitors are quiescent or proliferative and express nestin, Glial fibrillary acidic protein (GFAP), and SRY-box 2 (SOX2) (Seri et al., 2001; Moss et al., 2016). Type 1 cells extend radial processes through the GCL to the molecular layer (ML) and the hippocampal fissure. These cells also generate Type 2 progenitors, which express SOX2 and TBR2 and exhibit a multipolar morphology (Hutton and Pevny, 2011; Hodge et al., 2012). Type 2 progenitors undergo limited divisions and usually give rise to Type 3 progenitors (neuroblasts). Type 3 progenitors express doublecortin (DCX) and become dentate granule cells (GCs) (Martínez-Cerdeño and Noctor, 2018).

Additional cell types in the SGZ support neurogenesis. Adult-born GCs mature under gamma-aminobutyric acid (GABA) and glutamatergic inputs in the SGZ (Tozuka et al., 2005; Wang et al., 2005; Toni and Schinder, 2015). GABA influences intermediate neural progenitors and immature GCs in the dentate gyrus by promoting fate selection, proliferation, migration, and dendritic arbor maturation (Ge et al., 2006; Ge et al., 2007; Catavero et al., 2018). Tonic GABA promotes Type 1 progenitor quiescence (Song et al., 2012). Regulation of Type 1 progenitor proliferation occurs via parvalbumin (PV) interneurons, whereby PV activation represses proliferation and inactivation promotes proliferation (Song et al., 2013). In Type 2 progenitors, GABA signaling reduces proliferation and promotes their differentiation by inducing cell cycle exit (Deisseroth et al., 2004; Deisseroth and Malenka, 2005). Type 3 progenitors require synaptic GABAergic and glutamatergic inputs for their maturation and migration into deeper layers of the GCL (Kozareva et al., 2019).

Adult neurogenesis in the hippocampus is commonly dysregulated in temporal lobe epilepsy (TLE) (Kang et al., 2016). Prolonged seizure activity triggers a dramatic and transient rise in cell proliferation in the dentate gyrus (Parent et al., 1997; Gray and Sundstrom, 1998; Jessberger et al., 2005). Seizure activity particularly affects mitotically active progenitors and immature migrating DCX-expressing adult-born GCs (Parent et al., 1999; Hüttmann et al., 2003; Jessberger et al., 2005; Parent et al., 2006; Lugert et al., 2010). Even a single seizure-like discharge induces cell proliferation in the dentate gyrus (Bengzon et al., 1997; Römer et al., 2011). Seizure-generated proliferation of GCs appears to decline with increased seizure severity (Mohapel et al., 2004), due to inflammation (Ekdahl et al., 2003). Cell proliferation returns to baseline levels approximately 15 days to 4 weeks following the initial status epilepticus (SE) episode (Parent et al., 1997; Bonde et al., 2006; Römer et al., 2011), or may even be suppressed below baseline (Hattiangady et al., 2004; Kralic et al., 2005). Such changes may result from depletion of the pool of neural stem cells or altered environment in the SGZ (Jessberger and Parent, 2015). RGL cell polarity is altered by chronic TLE, assuming an inverted morphology, with radial processes directed toward the hilus of the dentate gyrus. This change in polarity may direct some RGL cells to transdifferentiate into astrocytes, suggesting a reduction in the neural stem cell pool that produces new GCs (Sasaki-Takahashi et al., 2020). A subset of GCs born after TLE induction in rodents develop abnormal structure, position, and orientation (Scharfman et al., 2007; Shapiro et al., 2008; Murphy et al., 2012). These hilar ectopic GCs migrate to an abnormal location-the hilus rather than the GCL. Hilar ectopic GCs form abnormal connections with neighboring GCs and CA3 pyramidal cells, forming a hyperexcitable dentate circuit that is more prone to spontaneous recurrent seizures (Scharfman et al., 2000; Scharfman, 2004; Scharfman and Bernstein, 2015; Althaus et al., 2019).

Additional structural changes observed in TLE include mossy fiber sprouting, enlarged GC dendritic arbor growth, reduced GC dendritic spines (Murphy et al., 2011), and loss of GABAergic interneurons in the hippocampus (Cossart et al., 2001). Seizure frequency significantly correlates with loss of GABAergic interneurons in or adjacent to the GCL (Buckmaster et al., 2017). Interneuron loss initially reduces GABAergic synapses on GCs, but later reaches control levels due to sprouting of the surviving interneurons (Thind et al., 2010).

Ablating adult-born GCs pre- or post-pilocarpine-induced SE reduces seizure burden (Cho et al., 2015; Hosford et al., 2016; Varma et al., 2019), but increases seizure duration (Hosford et al., 2016). This method significantly reduces seizures in mice with chronic epilepsy (Hosford et al., 2017). However, elimination of GCs born pre- or post-SE insult using methylazoxymethanol acetate reduces neurogenesis (ectopic and normotopic) but has no impact on seizure frequency (Zhu et al., 2017). Additionally, blocking post-SE neurogenesis using ganciclovir does not lead to long-term seizure suppression (Varma et al., 2019). This finding was also replicated in the kainic acid model of TLE, where ablation of adult-born GCs increased seizure severity (Iyengar et al., 2015). The mixed results of ablation studies in rodent epilepsy models may be due to the methods of ablation used and their differential effects on Type 1, 2, or 3 progenitors or the timing of the ablations. Yet, silencing adult-born GCs using the newer DREAD approach reduces seizures (Zhou et al., 2019; Lybrand et al., 2021). These findings underscore the role of aberrant adult neurogenesis in epileptogenesis and seizure development in rodent models of TLE (Cossart et al., 2001; Thind et al., 2010; Buckmaster et al., 2017).

Transplantation of mouse or human forebrain GABAergic interneuron progenitors is a powerful approach for reducing seizure burden in rodent models of TLE (Hunt et al., 2013; Cunningham et al., 2014; Henderson et al., 2014; Upadhya et al., 2019), suggesting that increasing GABAergic inhibition in the dentate gyrus may provide a mechanism for seizure suppression. Indeed, patch-clamp electrophysiological recordings in brain slices from mice with TLE showed increased spontaneous inhibitory postsynaptic currents in granule cells in slices with transplants (Henderson et al., 2014). The transplanted cells form extensive synaptic contacts associated with post-synaptic gephyrin puncta in the host brains and optogenetic stimulation of the transplanted cells induces strong inhibitory postsynaptic currents (Henderson et al., 2014; Gupta et al., 2019; Arshad et al., 2021). Despite mounting evidence that enhancing inhibitory synaptic transmission in the hippocampus or reducing adult neurogenesis are effective approaches for reducing seizures in rodent models of TLE, little is known about the underlying mechanisms for these therapeutic effects. Here, we addressed this gap in knowledge by comparing the effects of GABAergic progenitor transplantation on distinct stages of adult neurogenesis in naïve and epileptic mice.

## 2 MATERIALS AND METHODS

### Animals

Animal use conformed to protocols approved by the Wesleyan University Institutional Animal Care and Use Committee. Experiments were performed using group-housed C57BL/6NHsd mice (Table 1, Envigo) maintained in self-ventilating cages, under a 12 h light–dark cycle with light onset at 7 AM, room temperature of 25℃, humidity of 17%–20%, and light intensity of 193–215 lux at the cage level. Mice consumed food and water ad libitum. Mice were handled daily for 5 minutes for one week before inducing SE. Donor cells for GABAergic progenitor transplants were obtained on embryonic day 13.5 from VGAT-ChR2-eYFP^+^ embryos, by breeding VGAT-ChR2-eYFP transgenic males (Jackson Labs) with C57BL/6NHsd females (Envigo).

### Epilepsy model development

Pilocarpine was administered to induce SE in 5–6-week-old mice (Envigo, C57BL/6N strain). To generate mice with chronic TLE, we injected adult male or female mice at approximately 5–6 weeks of age and weighing 18–22 g with 0.07 mL methyl scopolamine (i.p., 0.5 mg/mL diluted in sterile saline; Sigma). After 30 min, we injected pilocarpine-HCl (i.p., 280 mg/kg), as described previously (Arshad and Naegele, 2020). Mice that did not develop SE were excluded from further study. SE was attenuated after 3 h, with systemic injection of midazolam (i.p., 0.04 mL; 9.1–11.1 mg/kg, Covetrus).

We used a multi-factorial design to compare the effects of interneuron transplantation vs. pilocarpine-induced SE on adult neurogenesis in a total of 99 (74 male and 25 female) mice. Forty-seven of 62 C57BL/6NHsd mice subjected to pilocarpine developed SE (47/62). Of these, 37 were male and 10 were female. Fifteen of 62 mice did not develop SE or died during pilocarpine induction (12 deaths and 3 failed to enter SE; those that failed to develop SE were not utilized further). Four SE mice were excluded due to damaged dentate gyri from unsuccessful targeting of stereotaxic injections. Of the 43 surviving SE mice, equivalent numbers were randomly assigned to SE-Sham or SE-Transplant (TX) groups; 22 received stereotaxic injections of fetal GABAergic progenitors into the dentate gyrus (SE-TX), and 21 received control stereotaxic injections of neural transplant media, without cells (SE-Sham). Of the 43 SE mice, 33 mice received systemic injections of BrdU, and 10 mice received systemic injections of EdU (see below).

An additional 36 age-matched control mice received an injection of methyl scopolamine followed by saline, instead of pilocarpine (and did not receive midazolam). These mice were designated “naïve” controls (26 males; 10 females), of which 12 received stereotaxic injections of fetal GABAergic progenitors into the dentate gyrus (Naïve-TX) and 24 received stereotaxic injections of transplant media without cells (Naïve-Sham). Of the control mice, 30/36 received BrdU and 6/36 received EdU (see below).

### Medial ganglion eminence (MGE) dissection, dissociation, and transplantation

Embryonic day 13.5 embryos were obtained by timed breeding of VGAT-ChR2-YFP adult male mice (stock number: 014548, The Jackson Labs) to wild-type females (C57Bl/6NHsd, Envigo). VGAT-ChR2-YFP+ embryos were removed and MGEs from each YFP+ embryo were isolated as described (Arshad et al., 2021). Following trypsin digestion and trituration to obtain a single-cell suspension, cells were re-suspended at a concentration of 1 × 10^5^ cells/μl in transplantation media (Leibovitz’s L-15 (Gibco, 11415-064), supplemented with caspase inhibitor (Promega G7231), B27 (Gibco, 17504-044), fibroblast growth factor (Sigma, F0291), and epidermal growth factor (Invitrogen, 53303-018) (Henderson et al., 2014). MGE-derived progenitors were transplanted, by stereotaxic injection, bilaterally into four sites in the hilus of the dentate gyrus (AP −2.5 mm, ML ±2.1 mm, DV −2.2 and −1.8 mm). Sham operates for SE and naïve control groups received 4 stereotaxic injections of the transplant media without MGE cells into four sites in the dentate gyrus with the same stereotaxic coordinates. Mice were returned to the home colony until further use. Naïve mice were group-housed (2–3 mice/cage) in the home colony; SE mice were individually-housed to prevent aggression.

### Immunohistochemistry

Mice were perfused with 4% paraformaldehyde in 0.1 M sodium phosphate buffer (pH 7.4), and brains were removed from the skull, postfixed for 15 h at 4°C, and equilibrated in ascending sucrose solutions (10%, 20%, and 30% sucrose in 0.1 M sodium phosphate buffer, pH 7.4). Forty-μm cryostat sections in the horizontal plane were collected in 0.1 M sodium phosphate buffer saline (PBS). Immunofluorescent staining to detect transplanted GABAergic interneurons expressing ChR2-EYFP was performed with chicken anti-GFP (GFP 1020, Aves), in combination with neurogenic markers. Type 1 progenitors were identified by immunostaining for GFAP (mouse mAb anti-GFAP; #IF03L, Millipore Sigma) and SOX2 (rabbit polyclonal anti-SOX2 #5603, Millipore Sigma). Type 2 progenitors were identified by positive immunoreactivity for SOX2 without co-expression of GFAP. Type 3 progenitors were identified by immunostaining with guinea pig anti-DCX (#2253, Millipore Sigma). Free-floating sections were blocked in 5% normal goat serum in PBS containing 0.3% Triton-X100 for one hour, then incubated for 18–22 h at RT while agitating in primary antibodies in PBS with 3% normal goat serum and 0.3% Triton-X100. Sections were washed in 0.1 M PBS, then incubated in goat anti-chicken IgY Alexa Fluor 488 (1:1000, #A11039, Life Technologies) and one or more of the following antibodies: goat anti-rabbit IgG Alexa Fluor 568 (1:1000, #A11036, Invitrogen), goat anti-mouse IgG Alexa Fluor 647 (1:1000, #A21242, Invitrogen), or goat anti-guinea pig IgG Alexa Fluor 647 (1:1000, #A21450, Invitrogen) for 1 h. The sections were rinsed, incubated in DAPI (1:2000, #62248, ThermoFisher Scientific), rinsed, and mounted on Superfrost Plus slides (#12-550-15, Fisher Scientific) in Prolong Diamond with DAPI (#P36966, ThermoFisher Scientific).

### BrdU labeling and detection

To examine effects of transplantation on cell proliferation, mice received a single dose of BrdU (i.p; 150 mg/kg; #B5002, Sigma-Aldrich), after survival periods of 1-, 2-, or 4-weeks following stereotaxic transplantation of MGE-derived progenitors or injections of media. Two hours after BrdU was injected, they were euthanized by injection of pentobarbital solution (0.1 mL i.p., Fatal Plus, Henry Schein, 035946) and we immediately performed transcardiac perfusion with ice-cold perfusion rinse buffer (0.1 M sodium phosphate buffer, pH 7.4) followed by 4% paraformaldehyde in 0.1 M sodium phosphate buffer (pH 7.4). To examine the effects of GABAergic progenitor transplantation on survival of adult-born GCs in the dentate gyrus, BrdU was injected one week following GABAergic progenitor transplantation, and the mice survived for an additional one or three weeks before they were euthanized and perfused, as described above. The brains were sectioned the day of staining. Sections were incubated in 1N HCl for 20 minutes at 37°C to denature the DNA and expose the incorporated BrdU, and the acid was neutralized by immersing sections in Trizma base (pH 8.5) for 2×5 min. Additional permeabilization was achieved by incubating sections in PBS with 1% Triton-X for 30 min. Next, sections were treated with anti-BrdU (Rat anti-BrdU, 1:500; MCA, 2060, BioRad) antibody solution followed by washes in PBS and secondary antibodies were applied. Nuclei were labeled with DAPI (1:2000, 62248, ThermoFisher Scientific), rinsed, and sections were mounted onto Superfrost Plus glass slides (#12-550-15, Fisher Scientific) in Prolong Diamond with DAPI (P36966, ThermoFisher Scientific).

### EdU labeling and detection

Due to loss of the DCX antigen associated with the HCL treatment during BrdU immunostaining described above, we performed additional proliferation experiments with 5-ethynyl-2′-deoxyuridine (EdU), an alternative thymidine analog, in combination with the Click-It fluorescent detection method. Two weeks after pilocarpine-induced SE, mice received i.p. injections of EdU (50 mg/kg) (#A10044, Invitrogen) to label proliferating cells. EdU was injected 4 times in one day, at two-hour intervals, over a period of 8 hours. Mice were perfused one week after EdU injections. Brains were removed and free-floating 40-μm cryostat sections were cut in the coronal plane. Sections were blocked with 5% normal goat serum and 0.3% PBST. Primary and secondary antibodies were diluted with PBST containing 3% NGS. Sections were incubated overnight at 4°C with guinea pig anti-doublecortin antibody (#AB2253, Millipore Sigma). After washing in PBS, sections were incubated for 1 h at room temperature with goat anti-guinea pig Alexa Fluor 647 secondary antibody (#A21450, Life Technologies). Sections were then incubated in the Click-IT reaction, according to the manufacturer’s instructions (#C10338, Life Technologies), to label incorporated EdU with Alexa Fluor 555. Sections were mounted on SuperFrost Plus slides in Prolong Diamond Antifade Mounting Medium.

### Quantitative confocal microscopy

To quantify immunofluorescently-labeled adult-born cells, we collected a 1:10 series of 40-μm frozen sections (400 μm apart), performed immunostaining, and obtained confocal images (Leica TCS SP8 confocal microscope) at 1-μm steps through the z-axis, with a Leica 20x/0.75 (zoom factor of 0.75) or 63x/1.4 objective. Counts of labeled neurons were made in 4 sections at 4 different levels along the dorso-ventral axis of the hippocampus. The region of interest (ROI) was defined within the dentate gyrus (hilus, GCL, SGZ) for counting labeled cells, avoiding guard zones, consisting of 5 μm at the top and bottom of each image. Within the ROI, counts were performed in optical sections spaced apart by 1-μm intervals; this was typically 20–22 optical slices along the z-plane of the 40-μm thick section. The SGZ was defined as the zone within one granule cell soma diameter (~10 μm) from the margin of the GCL (Varma et al., 2019). Hilar ectopic GCs were defined by their locations beneath the SGZ within 20 μm, or the equivalent of two cell bodies, away from the GC body layer (Singh et al., 2015).

### Quantitative analyses of granule cell migration

Migration of DCX^+^ cells into the GCL was assessed in flattened confocal images of brain sections ~ −2.36 mm ventral to the stereotaxic coordinates for the bregma. For each treatment group, the ROI was defined as a square box, measuring100 μm x 100 μm, located in the middle of the lateral blade of the dentate gyrus. The migration distance was measured from the center of the soma to the border of the GCL/SGZ, as described previously (Duveau et al., 2011).

### Statistical analysis

Values are expressed as mean ± standard error of the mean (SEM). For all comparisons, one-way ANOVA or t-tests were performed, followed by post hoc tests, whenever appropriate. Statistical tests were performed using GraphPad Prism, V7. The confidence interval was 95%. Differences were considered statistically significant at ∗p < 0.05, ∗∗p < 0.01, ∗∗∗p < 0.001, ****p < 0.0001.

## RESULTS

While previous work indicated that transplantation of GABAergic progenitors, or optogenetic stimulation of endogenous GABAergic interneurons are effective in suppressing seizures in mouse models of TLE, the mechanisms underlying these effects remain unclear. To investigate whether GABAergic progenitor transplantation corrects seizure-induced abnormalities in adult neurogenesis, we quantified the different populations of neural stem cells and progenitors in epileptic vs. naïve control mice, after immunostaining and performing high-resolution confocal microscopy.

### Does transplantation of GABAergic progenitors into the dentate gyrus alter the migration or location of adult born DCX^+^ cells in epileptic mice?

Prior work established that recurrent seizures in rodent TLE models are linked to increased numbers of adult born GCs that exhibit ectopic migration into the hilus (Scharfman et al., 2000; Scharfman, 2004; Jessberger et al., 2005; Parent et al., 2006; Scharfman et al., 2007; Danzer, 2018; Althaus et al., 2019). Therefore, we first investigated whether our seizure model in mice results in ectopic migration of immature GCs expressing DCX. To accomplish this, we compared DCX^+^ cells, in the absence of transplants, in epileptic mice (SE-Sham) vs. non-epileptic (naïve-Sham) mice (Fig.1 A-H). As expected, DCX^+^ cells were significantly increased in SE-Sham mice compared to naïve-Sham mice (Fig. 1 E-H). These effects were present throughout the dorso-ventral axis of the hippocampus but were more pronounced at dorsal levels (Supplementary Fig. 1). Strikingly, in mice with transplanted GABAergic progenitors, we observed a further significant increase in DCX^+^ cells in the GCL in both naïve and SE mice (Fig. 1 I-Q).

**Figure 1.**
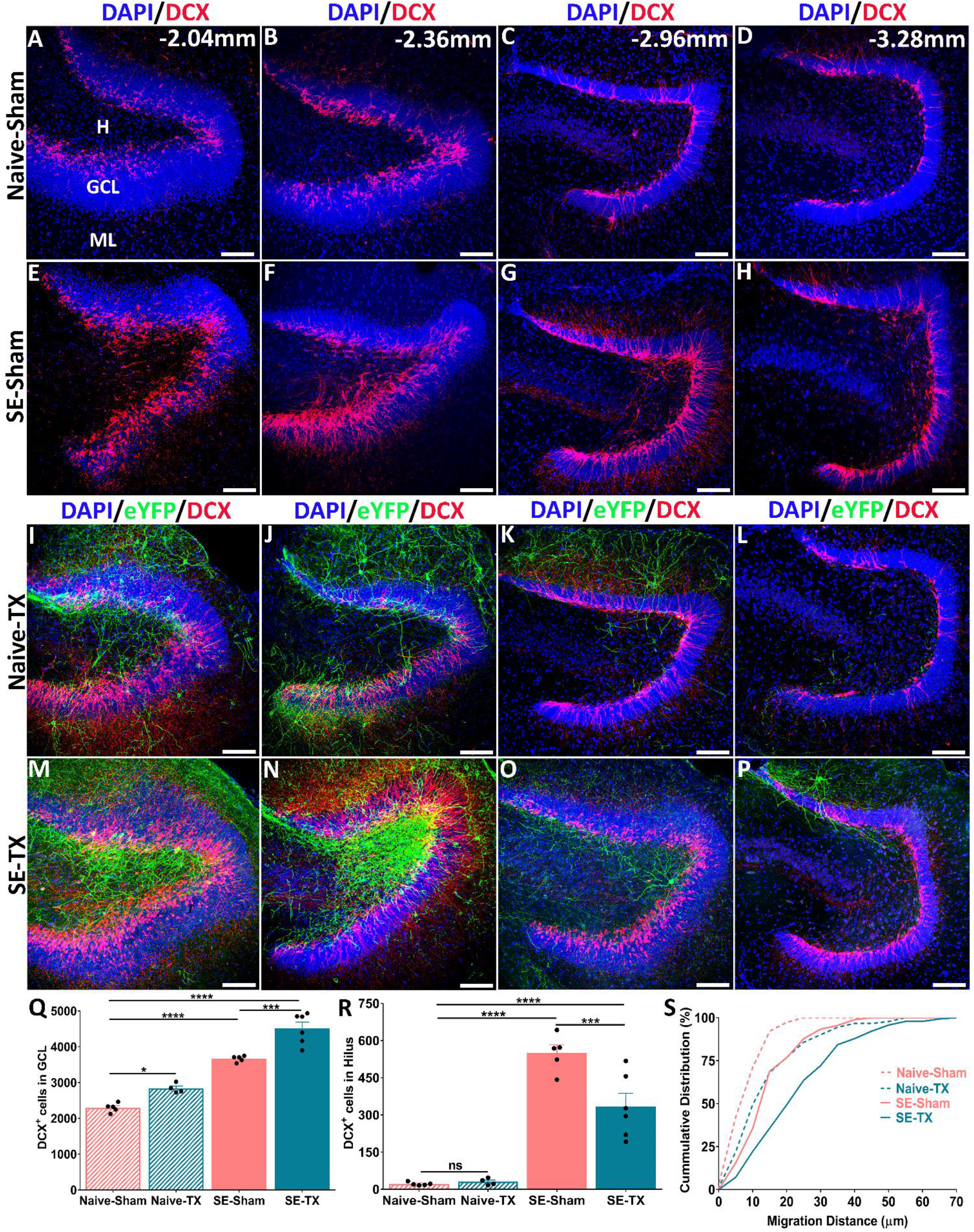
DCX^+^ adult-born granule cells in the dentate gyrus of the hippocampus increase in naïve and SE mice following transplantation of GABAergic cells. Naïve and SE mice that received injections of media (Sham; no cells transplanted) are shown (A-H) for comparison with naïve and SE mice with GABA progenitor transplants (I-E). A-D, Confocal images show DCX^+^ cells (red) in the DG of a naïve-Sham control mouse. Numbers in the upper right corner correspond to different dorsal-ventral levels from the bregma (FranklinKBJ, 2007). E-H, Confocal images of DCX^+^ cells (red) in an SE-Sham mouse. SE was associated with increased numbers of DCX^+^ cells and ectopic migration into the hilus of the dentate gyrus. I-P, Confocal images comparing DCX^+^ cells (red) at dorsal and ventral levels of the dentate gyrus in representative animals from the naïve-TX and SE-TX groups. Transplanted VGAT-ChR2-eYFP^+^ GABAergic progenitors (green) formed more extensive processes onto DCX^+^ cells (red) in more dorsal levels of the dentate gyrus. DAPI (blue) nuclear stain. Q, Quantification showed increased DCX^+^ cells in naïve mice with TX compared to naïve-Sham mice and significantly more DCX^+^ cells in SE-Sham operates compared to naïve-TX mice. SE-TX mice had significantly higher numbers of DCX^+^ cells compared to other groups. R, Quantification of the effects of GABAergic progenitor transplants on ectopic migration, showing that transplantation in SE mice resulted in significantly fewer ectopic DCX^+^ cells in the hilus of the dentate gyrus. S, Graphical representation of the cumulative distribution (%) of DCX^+^ cells in the four treatment groups based on the distance of radial migration (microns) from the SGZ. Significant differences were observed in the migration of DCX^+^ cells in the GCL. [n in each group: Naïve-Sham = 5; Naïve-TX = 4; SE-Sham = 5; SE-TX = 6. Statistics: (Q) ANOVA F(3,16) = 76.54; p<0.0001; Tukey’s multiple comparisons test DF=16; *Adjusted p-value (Naïve-Sham vs Naïve-TX) = 0.02; *** Adjusted p-value (Naïve-Sham vs SE-Sham) <0.0001; **** Adjusted p-value (Naïve-Sham vs SE-TX) < 0.0001; ***Adjusted p-value (SE-Sham vs SE-TX) = 0.0003); (R) ANOVA F(3,16) = 47.71; p<0.0001; Tukey’s multiple comparisons test DF= 16; ****Adjusted p-value (Naïve-Sham vs SE-Sham) <0.0001; ***Adjusted p-value (SE-Sham vs SE-TX) = 0.009; (S) Naive-Sham = 38 cells; Naïve-TX= 91 cells; SE-Sham = 90 cells; SE-TX= 141 cells; Kolmogorov-Smirnov test; *approximate p-value = 0.03 (Naïve-Sham vs Naïve TX); *approximate p-value = 0.01 (Naïve-Sham vs SE-Sham); **approximate p-value = 0.009 (Naïve-Sham vs SE-TX); ns approximate p-value = 0.3 (SE-Sham vs SE-TX)]. Scale bar = 100 μm.

These observations suggest that hilar GABAergic progenitor transplants may enhance radial migration or reduce ectopic migration in the DG of TLE mice. To distinguish between these alternatives, we compared numbers of hilar ectopic DCX^+^ cells in the four treatment groups of mice. As expected, Naïve-Sham mice had very few hilar ectopic DCX^+^ cells (Fig. 1R; naïve-Sham). In contrast, SE mice had many ectopic DCX^+^ cells (Fig. 1R; SE-Sham). Moreover, in SE mice, GABAergic progenitor transplants significantly reduced the number of ectopic cells (Fig. 1R; SE-TX). Taken together, these findings suggest that hilar transplants of GABAergic neurons inhibit ectopic migration of adult-born cells and promote migration into the GCL. To further investigate possible effects on migration, we next asked whether DCX^+^ cells migrated further distances into the GCL from the SGZ–GCL border in mice that received transplants by plotting the average migration distance for each group of mice (Duveau et al., 2011). As shown in Fig. 1S, DCX^+^ cells migrated significantly further in SE mice with transplants (SE-TX), compared to other groups (SE-Sham; naïve-Sham).

The increased migratory distances of adult-born granule neurons into the GCL and decreased ectopic migration into the hilus in SE-TX mice are consistent with previous work showing that GABAergic cells can influence the rates and distances that developing neurons migrate. However, prior work also supports a role for GABAergic inputs in survival of adult-born neurons in the dentate gyrus (Ge et al., 2006; Ge et al., 2007; Dieni et al., 2013; Song et al., 2013). Therefore, an alternative possibility is that GABAergic transplants enhance the survival of adult-born neurons that have successfully migrate into the GCL. To distinguish between these possibilities, we analyzed cell survival at different time points, after pulse-labeling proliferating cells with BrdU one week after transplanting GABAergic progenitors (Fig. 2A). Short-term survivals lasting two hours post-BrdU injection resulted in equivalent labeling of proliferative populations of cells in the SGZ of the dentate gyrus in naïve mice with or without transplants (Fig. 2B, C, D), suggesting that GABAergic transplants do not alter neural progenitor proliferation per se. We next compared GC survival at two different periods post-BrdU by comparing naïve mice with or without GABAergic progenitor transplants. After one- or three-week survival periods, the number of BrdU-labeled adult-born GCs in naïve mice with GABAergic progenitor transplants was significantly higher than in naïve mice without transplants (Fig 2E-G; H-J). These findings are consistent with the interpretation that transplants of GABA cells increase the survival of adult-born GCs. To determine whether these effects are local or systemic, we compared survival of BrdU-labeled cells at different dorso-ventral levels of the dentate gyrus of the hippocampus in naïve mice with (Naïve-TX) or without transplants (Naïve-Sham). BrdU-labeled new neurons were increased only at more dorsal levels of the hippocampus, where the transplanted GABAergic cells were located, suggesting a possible local effect on new neuron survival in mice with transplants (Supplementary Fig. 2E).

**Figure 2.**
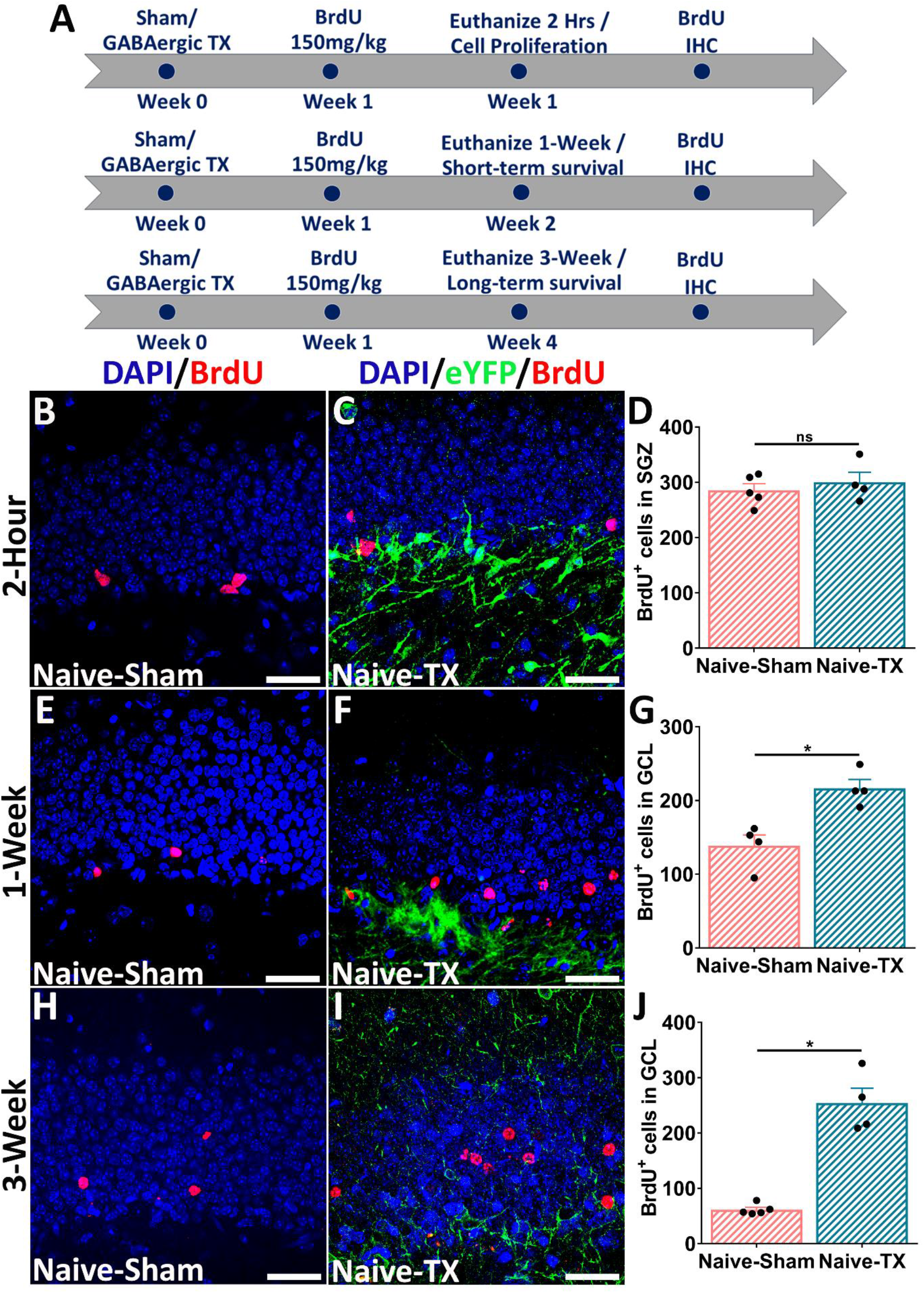
BrdU^+^ adult-born neurons increase following transplantation of GABAergic interneurons in naïve mice. A, Experimental timeline for study of cell proliferation and survival of adult-born neurons using the BrdU birthdating method. For the cell proliferation experiment, animals received a single dose of BrdU and were euthanized after 2 h. For the survival experiment, animals received a single dose of BrdU and were euthanized after 1 week or 3 weeks. B, Confocal image shows proliferating BrdU-labeled adult-born neurons (red) in the GC layer of the dentate gyrus from a naïve-Sham mouse. C, Confocal image shows proliferating BrdU-labeled adult-born neurons (red) in the GC layer of the dentate gyrus from a naïve transplant mouse. Transplanted VGAT-ChR2-eYFP^+^ GABAergic progenitors are shown in green. D, Quantification of the effects of GABAergic progenitor transplants on cell proliferation shows no difference in the number of proliferating BrdU-labeled adult-born neurons between the two groups. E, Confocal image shows 1-week-old BrdU-labeled adult-born neurons (red) in the GC layer of the dentate gyrus from a naïve-Sham mouse. F, Confocal image shows 1-week-old BrdU-labeled adult-born neurons (red) in the GC layer of the dentate gyrus from a naïve-TX mouse. Transplanted VGAT-ChR2-eYFP^+^ GABAergic progenitors are shown in green. G, Quantification shows a significant increase in the number of surviving 1-week-old BrdU-labeled adult-born neurons in naïve-TX mice vs naïve-Sham mice. H, Confocal image shows 3-week-old BrdU-labeled adult-born neurons (red) in the GC layer of the dentate gyrus from a naïve-TX mouse. I, Confocal images show 3-week-old BrdU-labeled adult-born neurons (red) in the GC layer of the dentate gyrus from a naïve-TX mouse. Transplanted VGAT-ChR2-eYFP^+^ GABAergic progenitors are shown in green. J, Quantification shows a significant increase in the number of surviving 3-week-old BrdU-labeled adult-born neurons in naïve-TX mice vs naïve-Sham mice. [n in each group: Naive-Sham= 5; Naive-TX= 4 in “2-Hour” group; Naive-Sham= 4; Naive-TX= 4 in “1-week” group; Naive-Sham= 5; Naive-TX= 4 in “3-week” group; Statistics: (G) Mann Whitney test (Naïve-Sham vs Naïve-TX) * Exact p-value= 0.02; (J) Mann Whitney test (Naïve-Sham vs Naïve TX) * Exact p-value = 0.01]. Scale bar = 25 μm.

We next performed similar experiments in SE mice (Fig. 3A) and found that GABAergic transplantation was not associated with an increase in proliferating cells in the SGZ at two-hours post injection (Fig. 3B-D) but was associated with increased survival of BrdU^+^ cells in the GCL three weeks post-injection (Fig. 3E-G). This effect of the transplants on survival of adult born BrdU-labeled cells at three weeks post-BrdU was also not distributed along the entire dorso-ventral axis of the hippocampus; BrdU-labeled cells were only significantly increased at more dorsal levels of the hippocampus, where the transplants were located, again suggesting a localized effect of the transplants on the survival of adult-born GCs (Supplementary Fig. 2F). To further confirm that these effects were due to survival of new neurons, rather than astrogliosis, we pulse labeled new cells with the thymidine analog EdU and performed dual staining for EdU incorporation and expression of DCX. The results show that that most, if not all EdU^+^ cells are also DCX^+^ (Supplementary Fig. 3). Thus, GABAergic progenitor transplantation is linked to significantly increased survival of adult-born GCs and further migration into the GCL, and these effects are present in naïve and epileptic mice.

**Figure 3.**
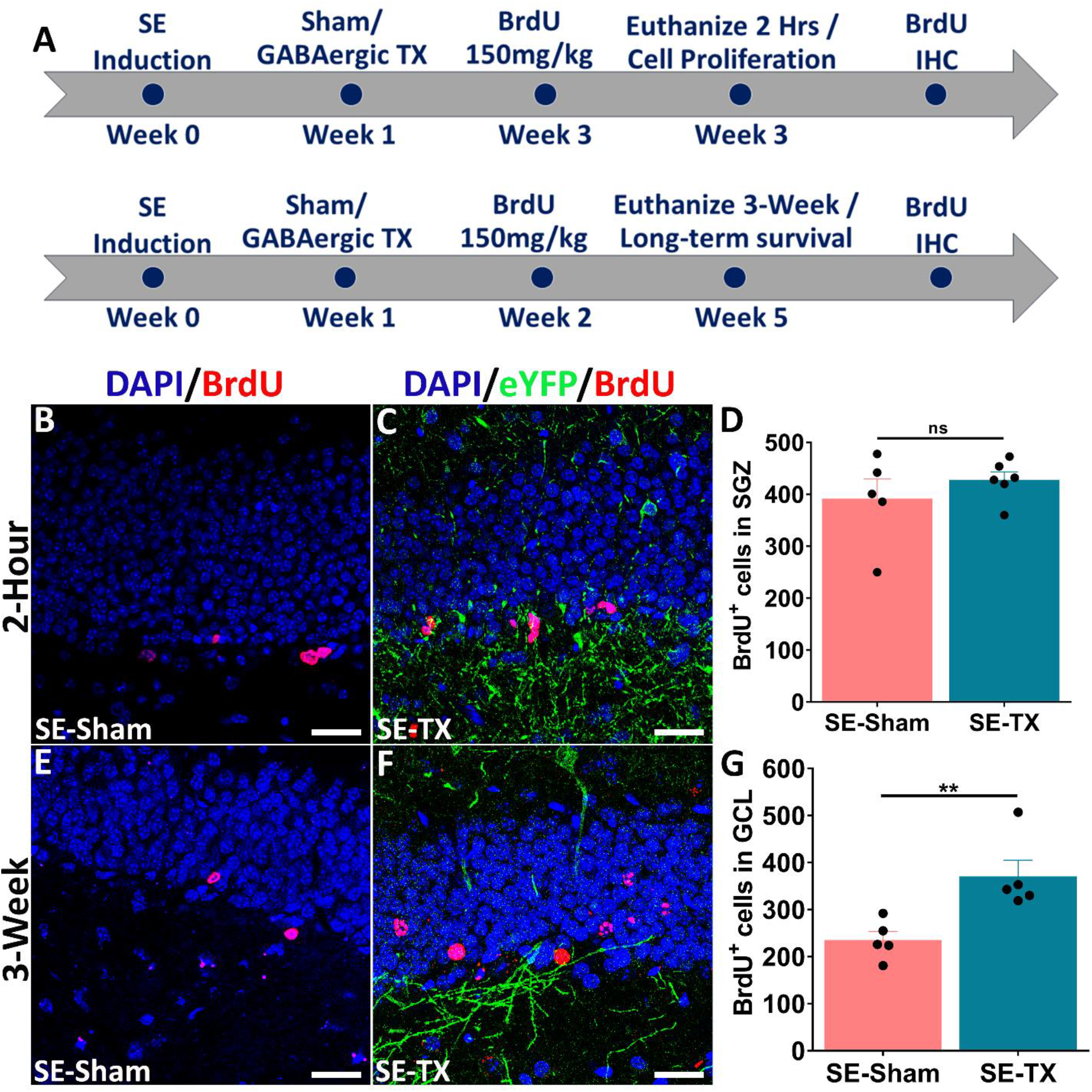
Transplanted GABAergic progenitors increase survival of adult-born neurons in SE mice. A, Experimental timeline for study of proliferation and survival of adult-born neurons using the BrdU birthdating method. For the cell proliferation experiment, animals received a single dose of BrdU and were euthanized after 2 h. For the survival experiment, animals received a single dose of BrdU and were euthanized after 3 weeks. B) Confocal image shows proliferating BrdU-labeled adult-born neurons (red) in the GC layer of the dentate gyrus from an SE-TX mouse. Transplanted VGAT-ChR2-eYFP^+^ GABAergic progenitors are shown in green. D, Quantification shows no difference in the number of proliferating BrdU-labeled adult-born neurons between the two groups. E, Confocal image shows 3-week-old BrdU-labeled adult-born neurons (red) in the GC layer of the dentate gyrus from an SE-Sham mouse. F, Confocal image shows 3-week-old BrdU-labeled adult-born neurons (red) in the GC layer of the dentate gyrus from an SE transplant mouse. Transplanted VGAT-ChR2-eYFP^+^ GABAergic progenitors are shown in green. G, Quantification shows a significant increase in the number of surviving 3-week-old BrdU-labeled adult-born neurons in SE transplant mice vs SE sham mice. [n in each group: SE-Sham= 5; SE-TX= 6 in “2-Hour” group; SE-Sham= 5; SE-TX= 5 in “3-week” group; Statistics: (G) Mann Whitney test (SE-Sham vs SE-TX) * Exact p-value= 0.007]. Scale bar = 25 μm.

### Are transplants of GABAergic neurons associated with alterations in other neural progenitor populations in the SGZ?

Type 1 progenitors are specialized RGL cells that give rise to Type 2 progenitors and self-renew. Optogenetic stimulation of endogenous dentate GABAergic interneurons promotes quiescence of Type 1 progenitors (Bao et al., 2017), suggesting that some of the effects we observed might be a consequence of the transplants inducing quiescence of Type 1 progenitors. To address this question, we next asked whether transplantation of GABAergic neurons alters the proportion of Type 1 progenitors (GFAP^+^/SOX2+) (Fig. 4A-C). Comparison of sham operates of naïve and epileptic mice showed that Type 1 progenitors were significantly reduced in mice with SE, consistent with prior studies (Singh et al., 2015; Sasaki-Takahashi et al., 2020); GABA progenitor transplantation did not rescue Type 1 progenitors (Fig. 4D).

**Figure 4.**
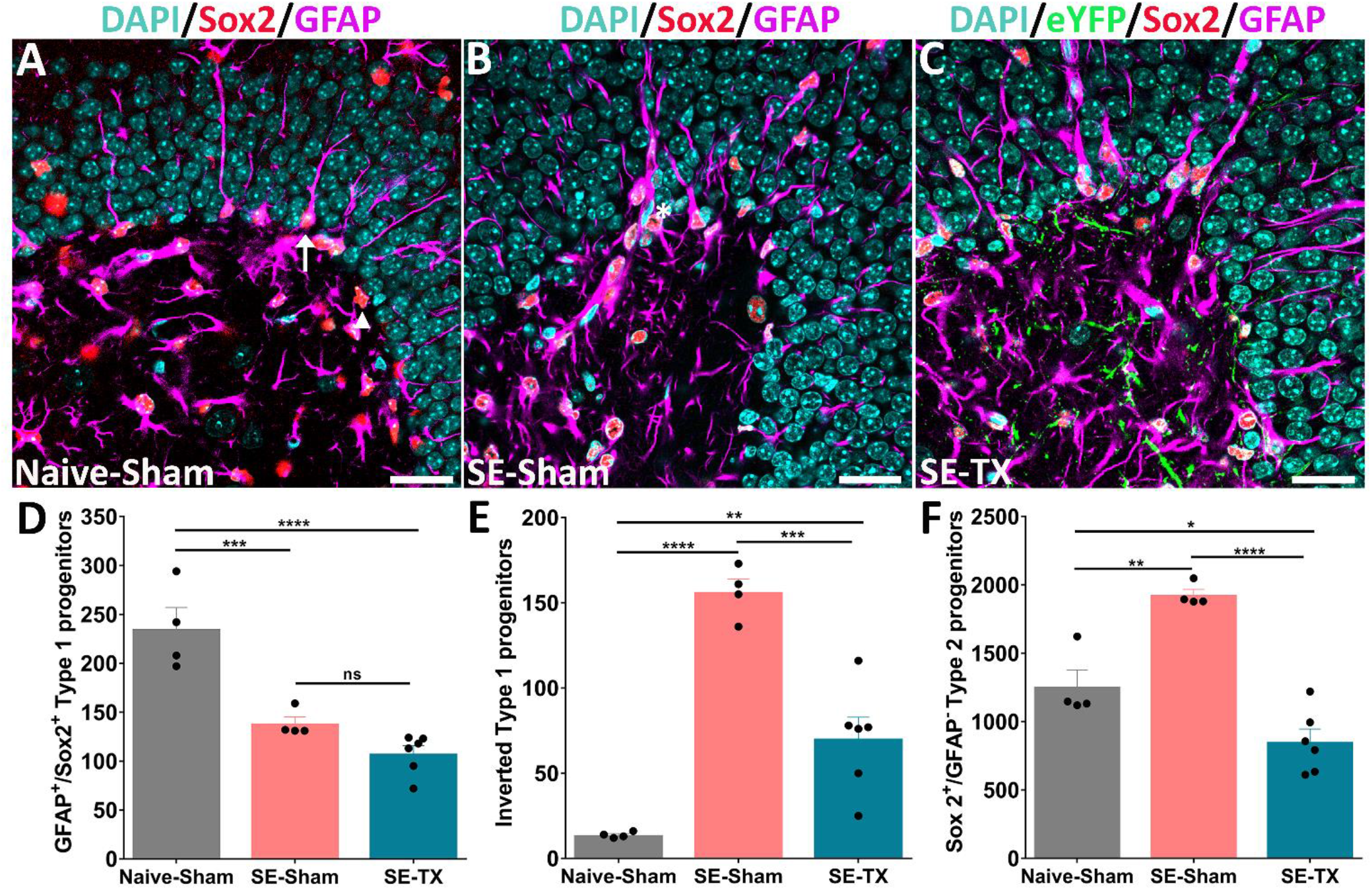
Type 2 and inverted Type 1 progenitors are reduced following transplantation of GABAergic progenitors. A-C, Confocal images show Type 1(GFAP^+^/ SOX2^+^) and Type 2 (SOX2^+^/ GFAP^**-**^) progenitors in the SGZ of the dentate gyrus in naïve-Sham, SE-Sham, and SE-TX groups respectively. The white arrow is pointing toward a Type 1 progenitor located in the SGZ of the dentate gyrus, with its apical process extending through the granular cell layer towards the pia and the hippocampal fissure as shown in panel A. In the case of severe SE, the orientation of the apical process is inverted and is extended towards the hilus as shown by an asterisk (white) in panel B. Type 2 progenitors express SOX2 and reside in the SGZ of the dentate gyrus, indicated by a white arrowhead as shown in panel A. GFAP-positive cells are shown in purple while SOX2-positive cells are shown in red. DAPI is used as a counterstain (cyan) and transplanted GABAergic progenitors are shown in green. D, Graph showing a significant reduction in the number of Type 1 progenitors in the SGZ of the dentate gyrus in SE mice with or without GABAergic transplants compared to naïve-Sham mice. E, Quantification shows fewer inverted Type 1 progenitors in SE mice following transplantation of the GABAergic progenitors vs SE-Sham mice. F, Graph shows a significant reduction in the number of SOX2^+^/GFAP^**-**^type 2 progenitors in the SGZ of the dentate gyrus in SE-TX vs SE-Sham mice. [n in each group: Naive-Sham =4; SE-Sham = 4; SE-TX= 6; Statistics: (D) ANOVA F_(2,11)_ = 26.59; p<0.0001; Tukey’s multiple comparisons test DF=11; ***Adjusted p-value (Naïve-Sham vs SE-Sham) = 0.008; **** Adjusted p-value (Naïve-Sham vs SE-TX) <0.0001; (E) ANOVA F_(2,11)_ = 42.02; p<0.0001; Tukey’s multiple comparisons test DF= 11; ****Adjusted p-value (Naïve-Sham vs SE-Sham) = 0.0059;****Adjusted p-value (SE-Sham vs SE-TX) < 0.0001; F) ANOVA F(2,11) = 32.72; p<0.0001; Tukey’s multiple comparisons test DF= 11; **Adjusted p-value (Naïve-Sham vs SE-Sham) = 0.002;*Adjusted p-value (Naive-Sham vs SE-TX) < 0.0001; ****Adjusted p-value (SE-Sham vs SE-TX) < 0.0001]. Scale bar = 25 μm.

The radial glial processes from Type 1 progenitors normally extend through the GCL into the inner third of the molecular layer and contact blood vessels. These processes provide structural support for radial migration of adult-born neurons and appear to regulate synapse formation between adult-born GCs and afferent inputs (Xu et al., 2015; Moss et al., 2016). In epilepsy, inversion of the radial glial progenitors may be associated with ectopic migration of adult-born GCs, as the Type 1 cells may adopt an inverted morphology (Fig. 4B), with processes extending toward the hilus (Sasaki-Takahashi et al., 2020). Given our prior results showing many ectopic DCX^+^ cells in SE mice, we next quantified abnormal GFAP^+^/SOX2^+^ Type 1 progenitors in the SGZ with inverted processes that extended into the hilus. While we found that these cells were significantly increased in SE mice, they were reduced in SE mice with transplants (Fig. 4E). Additionally, there was a significant reduction in the number of Type 2 (GFAP^-^/SOX2^+^) progenitors in the SGZ of the dentate gyrus in SE mice with GABAergic transplants (Fig. 4F).

### Does transplantation of GABAergic progenitors alter the proliferation of Type 1 progenitors in the SGZ of the dentate gyrus in SE mice?

To determine whether GABA progenitor transplantation affects Type 1 progenitor proliferation in SE mice, we examined mitotically active progenitors with BrdU pulse-labeling and performed immunostaining to identify Type 1 RGL progenitors by expression of GFAP (GFAP^+^/BrdU^+^) (Fig. 5A). Confirming our earlier studies of proliferation, we observed no significant differences in the number of dividing Type 1 progenitors in the SGZ of the dentate gyrus between the two groups of epileptic mice (SE-Sham vs SE-TX) (Fig. 5B-I).

**Figure 5.**
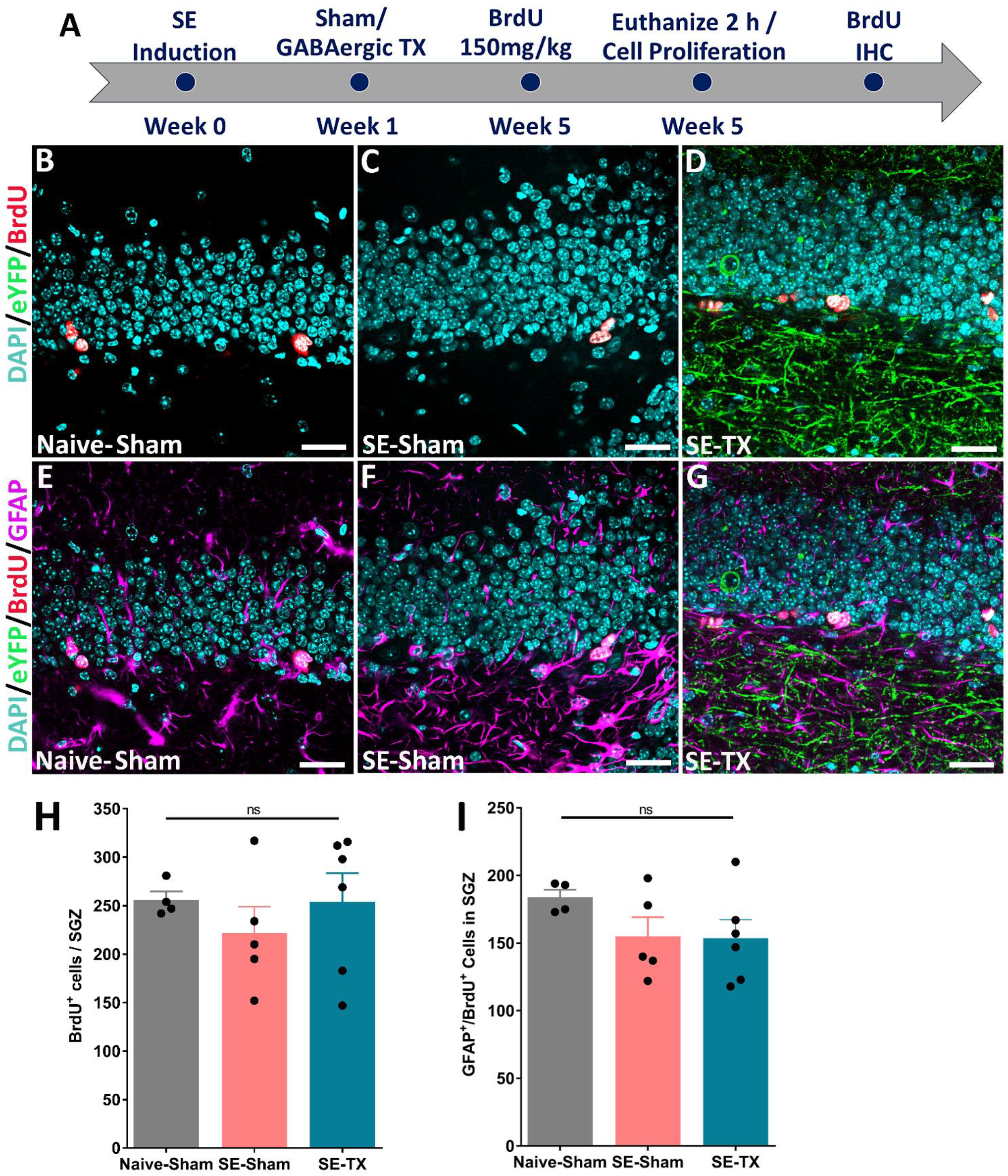
GABAergic progenitor transplant does not affect proliferation of Type 1 progenitors in the SGZ of the dentate gyrus of epileptic mice. A, Experimental timeline, mice received BrdU injection 5 weeks after SE induction and were perfused two hours after BrdU injection. B-D) Confocal images show proliferating BrdU-labeled adult-born neurons in the SGZ of the naïve-Sham, SE-Sham, and SE-TX mice. H, Quantification shows no significant difference in the number of BrdU^+^ cells across all groups. E-G) Confocal images show proliferating Type 1 progenitors (GFAP^+^/ BrdU^+^) in the SGZ of the naïve-Sham, SE-Sham, and SE-TX mice. I, Graph shows no significant difference in the number of proliferating Type 1 progenitors (GFAP^+^/ BrdU^+^) across all three groups. [n in each group: Naive-Sham =4; SE-Sham = 5; SE-TX= 6; Statistics: (H) ANOVA F(2,12) = 0.5392; p = 0.602; Tukey’s multiple comparisons test DF=12; ns Adjusted p-value (Naïve-Sham vs SE-Sham) = 0.66; ns Adjusted p-value (Naïve-Sham vs SE-TX) = 0.998; ns Adjusted p-value (SE-Sham vs SE-TX) = 0.6432 (I) ANOVA F(2,12) = 1.529; p = 0.2561; Tukey’s multiple comparisons test DF=12; ns Adjusted p-value (Naïve-Sham vs SE-Sham) = 0.3328; ns Adjusted p-value (Naïve-Sham vs SE-TX) = 0.2779; ns Adjusted p-value (SE-Sham vs SE-TX) = 0.9968]. Scale bar = 25 μm.

## 4 DISCUSSION

The major findings of this study in adult mice with TLE are that GABAergic progenitor transplants in the dentate gyrus of the hippocampus enhance survival and migration of adult-born GCs. Both SE and naïve mice receiving GABAergic progenitor transplants have significant increases in surviving adult-born GCs and these newborn cells exhibit enhanced migration from the SGZ into the GCL. Further, transplanted SE mice have a depletion in the number of inverted Type 1 and Type 2 progenitors in the SGZ. These results support that intrahippocampal transplants of GABAergic progenitors promote survival of adult-born neurons, possibly through accelerated Type 2 progenitor differentiation into DCX^+^ immature neurons, reduced programmed cell death, or both mechanisms.

### Transplantation of GABAergic progenitors enhances survival of adult-born granule cells

In rodents, 30–80% of adult-born GCs in the hippocampus undergo programmed cell death (Cameron and McKay, 2001; Dayer et al., 2003; Sun et al., 2004; Kuhn, 2015; Denoth-Lippuner and Jessberger, 2021). Moreover, endogenous populations of hippocampal GABAergic interneurons promote survival of dentate GCs (Dieni et al., 2013), while endogenous subsets of PV-expressing hippocampal GABAergic interneurons specifically enhance survival of adult-born GCs (Song et al., 2013). We previously found that E13.5 MGE-derived transplants consist of approximately 37% PV+ GABAergic interneurons; many of the transplanted cells distribute throughout the molecular and GCL layers, where upon maturation, they form extensive inhibitory synaptic contacts to the dendrites of adult-born GCs (Hunt et al., 2013; Henderson et al., 2014; Gupta et al., 2019). Therefore, the transplanted cells may be responsible for increasing PV+ synaptic contacts with adult-born GCs, contributing to their survival and integration into synaptic networks.

The cAMP response element-binding protein (CREB) is a downstream effector of GABA signaling and its activated form, phospho-CREB, is expressed by a high percentage of adult-born GCs (Nakagawa et al., 2002; Redmond et al., 2002). Phosphorylation of CREB coincides with a developmental phase when adult-born neurons depolarize in response to GABA-signaling (Ge et al., 2007; Jagasia et al., 2009), which is essential for maturation of adult-born GCs (Wang et al., 2003; Ge et al., 2007; Ben-Ari et al., 2012). The depolarizing responses of adult-born GCs to GABA depends upon expression of NKCC1, as knockdown of NKCC1 in adult-born GCs decreases pCREB, impairs outgrowth of dendrites, and decreases GC survival (Ge et al., 2007; Jagasia et al., 2009). These studies suggest that transplants may increase GABA-induced depolarization, leading to enhanced survival through activation of CREB-signaling.

### Enhanced radial migration of adult-born granule cells into the GCL due to GABAergic progenitor transplants

We further observed an increase in the migration of DCX-expressing adult-born GCs into the GCL of the dentate gyrus following transplantation of GABAergic progenitors. Reelin is a large extracellular matrix glycoprotein expressed by GABAergic basket cells in the dentate gyrus of the hippocampus (Curran and D’Arcangelo, 1998; Pesold et al., 1998) that acts as a repellant signal during the migration of adult-born dentate GCs. Reelin exerts its effect by acting on a migration scaffold formed by Type 1 progenitors (Frotscher et al., 2003), directing the migration of adult-born GCs into the GCL (Wang et al., 2018). Loss of reelin+ cells in the hilus and molecular layer after epileptic seizures was linked to ectopic migration of adult-born GCs (Parent et al., 2006; Gong et al., 2007), leading to the suggestion that reelin deficiency in TLE causes a disruption in the normal pattern of radial migration of newborn neurons or neuroblasts (Orcinha et al., 2016; Orcinha et al., 2021).

We found a correlation between reduced numbers of inverted Type 1 progenitors and increased migration and survival of adult-born GCs in TLE mice with transplants. While additional studies are necessary to determine the precise molecular and cellular mechanisms for increasing survival and normalizing the direction of migration of adult-born GCs in TLE mice with GABAergic transplants, our findings suggest that the exogenous GABAergic cells may provide the signals for these phenomena. In agreement with prior studies, we found that mice with pilocarpine-induced epilepsy had reduced numbers of normal Type 1 progenitors with radial processes into the GCL (Sierra et al., 2015; Singh et al., 2015; Sasaki-Takahashi et al., 2020) but increased inverted Type 1 progenitors. While the prevalence of Type 1 cells with radial processes did not appear to be altered following GABAergic progenitor transplantation, inverted Type 1 progenitors were markedly reduced following transplantation. Inversion of Type 1 progenitor processes can occur due to re-orientation of the centrosome away from the GCL toward the hilus, and centrosomal reorientation can be induced by seizures (Sasaki-Takahashi et al., 2020). This change in orientation of Type 1 progenitors might be regulated by reelin, which modulates neuronal migration (Zhao et al., 2004). In the absence of Reelin signaling, Reelin treatment in hippocampal slice culture experiments was shown to rescue the correct radial orientation of Type 1 progenitors (Jossin, 2020). Thus, deficient reelin signaling in TLE may result in development of inverted morphology of Type 1 progenitors and contribute to ectopic migration of adult-born DCX^+^ cells in the hilus. Our findings suggest that transplants help to maintain normal radial fiber orientation and prevent ectopic migration of new GCs.

### A role of GABAergic transplants in regulating Type 2 progenitors

Seizures increase proliferation of Type 2 progenitors (Indulekha et al., 2010; Jiruska et al., 2013; Jessberger and Parent, 2015; Fu et al., 2019). Accordingly, our experiments showed that SE mice that received sham injections of media also had significantly increased Type 2 cells. Our findings extend prior work by showing that Type 2 cells in epileptic mice were reduced following GABAergic progenitor transplantation. The reduced number of SOX2-expressing Type 2 progenitors could be due to increased GABA signaling, causing more rapid differentiation into DCX-expressing GCs. In support of this interpretation, prior work showed that GABA receptor agonist treatment induces rapid differentiation of Type 2 progenitors due to downregulation of anti-neuronal genes and upregulation of pro-neural gene neuroD, required for differentiation of Type 2 progenitors into post-mitotic cells (Deisseroth et al., 2004; Deisseroth and Malenka, 2005; Tozuka et al., 2005).

In conclusion, the current study builds on our prior work establishing that GABAergic progenitor transplants reduce seizure burden in TLE mice. Here, we provide new insights on a possible mechanism of GABAergic progenitor transplant-mediated seizure suppression. Our findings suggest that GABAergic progenitor transplants enhance the survival of adult-born neurons in naïve and epileptic mice. GABAergic progenitor transplants in TLE mice may prevent hyperexcitability in the dentate circuit by reducing the number of ectopic adult-born GCs in the hilus, as most of these adult-born neurons migrate normally to integrate into the dentate GCL. Finally, we found a significant decrease in the number of inverted Type 1 progenitors in TLE mice with GABAergic progenitor transplants, suggesting disease-modifying effects of the GABAergic graft. Taken together, our findings indicate that GABAergic progenitor transplant-mediated seizure suppression may be due in part to restoration of normal patterns of neurogenesis and migration of adult born granule cells in TLE mice.

## FIGURE CAPTIONS

**Supplementary Figure 1.**
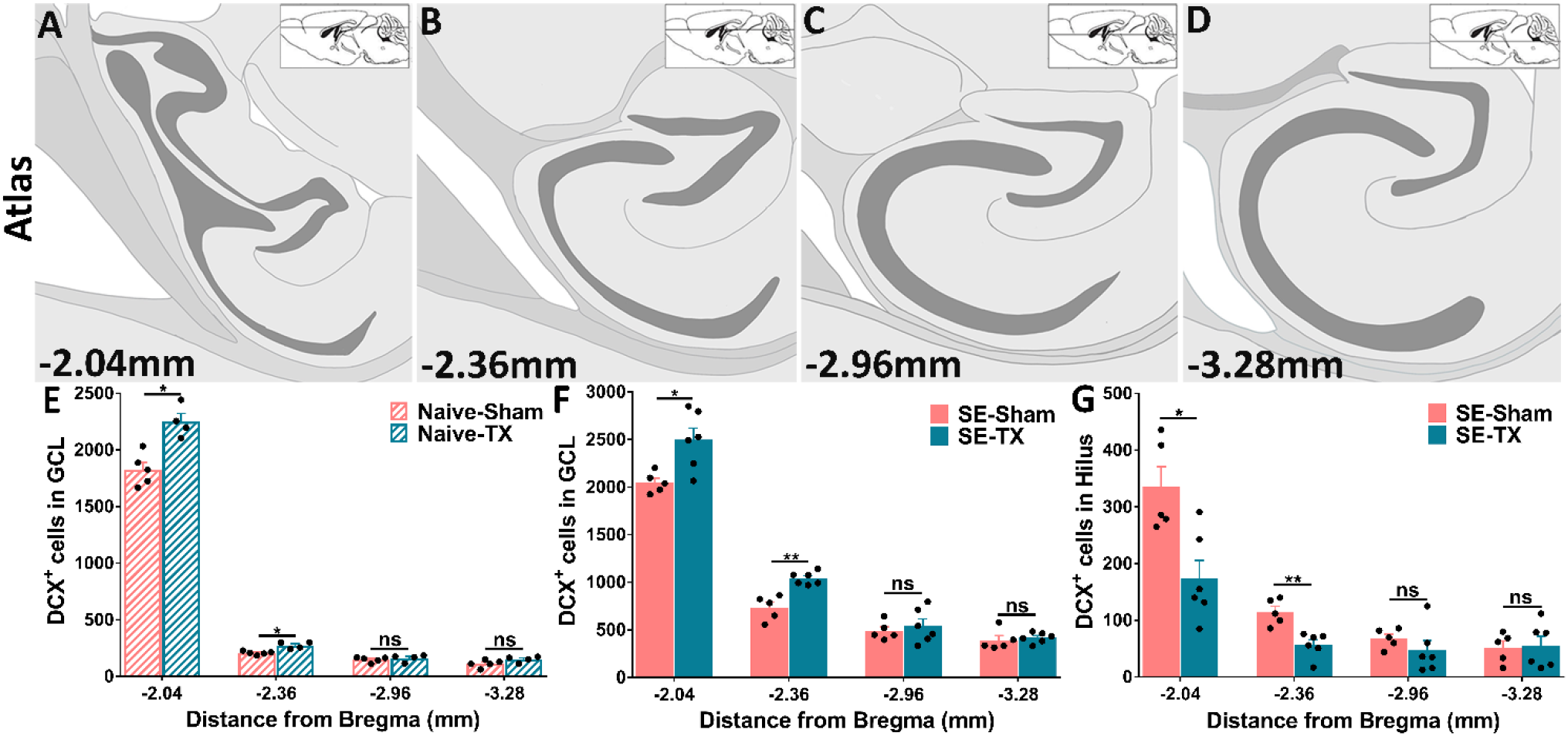
DCX^+^ adult-born neurons are increased in the dorsal hippocampus following transplantation of GABAergic interneurons. A-D, Modified atlas (FranklinKBJ, 2007) images showing an overview of the horizontal sections (distance from the bregma). E, GABAergic interneuron transplantation significantly increased the number of DCX^+^ cells in the dorsal hippocampus but not in the ventral hippocampus in naïve-TX mice compared to naïve-Sham mice. F, GABAergic interneuron transplantation significantly increased the number of DCX^+^ cells in the dorsal hippocampus but not in the ventral hippocampus in SE-TX mice compared to SE-Sham mice. G) Graph comparing the dorso-ventral distributions of DCX^+^ cells in the dentate gyrus shows the most significant reductions in ectopic adult-born GCs were in the more dorsal levels of the hippocampus. [n in each group: Naïve-Sham =5; Naïve-TX=4; SE-Sham = 5; SE-TX= 6; Statistics: (E) t-test, Multiple comparisons followed by the Holm-Sidak method (Naïve-Sham vs Naïve-TX) *Adjusted p-value = 0.012; (F) t-test, Multiple comparisons (SE-Sham vs SE-TX) *Adjusted p-value = 0.03; ** Adjusted p-value = 0.002; (G) t-test, Multiple comparisons (SE-Sham vs SE-TX) *Adjusted p-value = 0.02; **Adjusted p-value = 0.008]

**Supplementary Figure 2.**
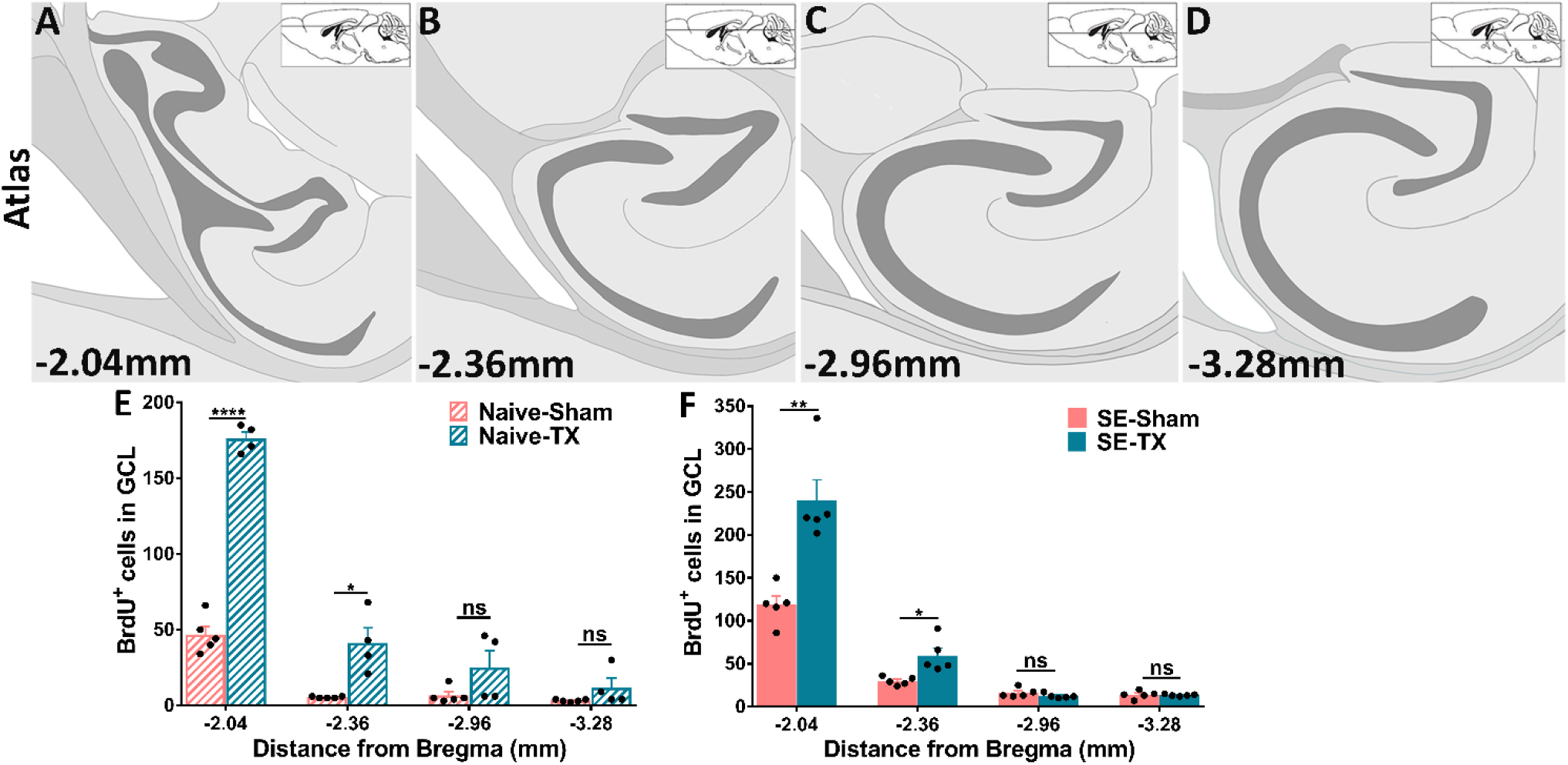
BrdU^+^ adult-born neurons are increased in the dorsal hippocampus following transplantation of GABAergic interneurons. A-D, Horizontal planes through the hippocampus modified from the mouse brain atlas (FranklinKBJ, 2007) showing dorsal to ventral levels through the dentate gyrus. E, A significant increase in the number of BrdU^+^ cells was found at more dorsal levels of the dentate gyrus in the naïve transplant group compared to the naïve-Sham group, corresponding to the levels of the dentate gyrus that contained the most transplant-derived GABAergic interneurons. F, A significant increase in the number of BrdU^+^ cells was found at more dorsal levels of the dentate gyrus in the SE-TX group vs. SE-Sham group. [n in each group: Naive-Sham =5; Naïve-TX=4; SE-Sham = 5; SE-TX= 6; Statistics: (E) t-test, Multiple comparisons followed by the Bonferroni-Dunn method (Naïve-Sham vs Naïve-TX) *Adjusted p-value = 0.018; **** Adjusted p-value = 0.00001 (F) t-test, Multiple comparisons (SE-Sham vs SE-TX) *Adjusted p-value = 0.04; ** Adjusted p-value = 0.0069]

**Supplementary Figure 3.**
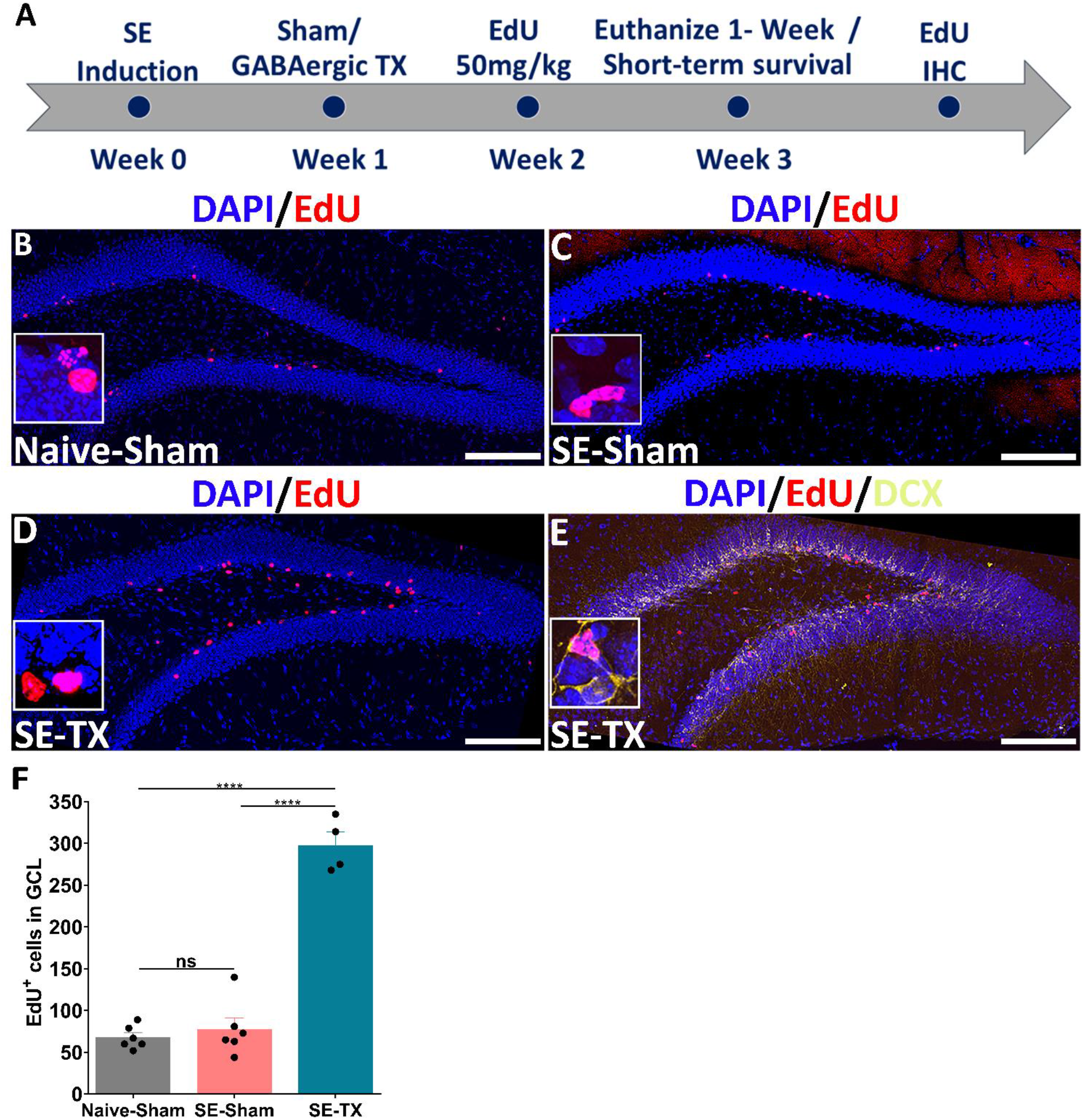
Transplantation of GABAergic progenitors significantly increases EdU^+^ adult-born neurons, confirming BrdU data. A, Experimental timeline to study adult-born neuron survival using the BrdU birthdating method. One week after receiving GABAergic transplants, mice received EdU (50 mg/kg) and were euthanized 1 week later. B, Confocal image shows 1-week-old EdU-labeled adult-born neurons (red) in the GC layer of the dentate gyrus from a naïve-Sham mouse. C, Confocal image shows 1-week-old EdU-labeled adult-born neurons (red) in the GC layer of the dentate gyrus from an SE-Sham mouse. D, Confocal image shows 1-week-old EdU-labeled adult-born neurons (red) in the GC layer of the dentate gyrus from an SE-TX mouse. DAPI (blue) was used as a counter stain. E, Confocal image shows 1-week-old EdU/DCX (golden) co-labeled adult-born neurons in the GC layer of the dentate gyrus from an SE-TX mouse. DAPI (blue) was used as a counter stain. Insets are magnified views of EdU^+^ cells. Bottom right inset, panel C, EdU^+^ cell co-labeled with DCX. G, Quantification shows a significant increase in the number of surviving 1-week-old EdU-labeled adult-born neurons in SE-TX mice vs SE-Sham mice. [n in each group: Naive-Sham =6; SE-Sham = 6; SE-TX= 4; Statistics: (F) ANOVA F(2,13) = 105.1; p <0.0001; Tukey’s multiple comparisons test DF=13; ns Adjusted p-value (Naïve-Sham vs SE-Sham) = 0.85; ****Adjusted p-value (Naïve-Sham vs SE-TX) < 0.0001; **** Adjusted p-value (SE-Sham vs SE-TX) < 0.0001]. Scale bar: 100 μm.

## Data Availability Statement

The raw data of this article will be made available by authors without undue reservation.

## Ethics Statement

The study was approved by the Wesleyan University Institutional Animal Care and Committee.

## Author Contributions

MNA and JRN designed the study. MNA and JRN performed all the experiments. MNA, SO, JJ, and BB analyzed the data. MNA and JRN wrote the manuscript with input from all the authors.

## Funding

This work was supported by the National Institute of Neurological Disorders and Stroke Grant R15NS072879-01A1, a Connecticut Stem Cell Established Investigator Grant, a Challenge Award from Citizens United for Research in Epilepsy (J.R.N), and grants in support of scholarship from Wesleyan University.

## Conflict of Interest

The authors declare that there are no commercial or financial relationships that could be construed as a potential conflict of interest.

## Acknowledgements

We would like to thank the former and current Naegele lab members Nicolas Cimino, Alina Widmann, Fitzroy Wickham, Marion Humphreys, Mitzi Adler-Wachter, Spencer Fox, Sophia Marra, and Calista Stevens. We thank Jeff Gilarde for technical assistance with confocal microscopy and Sera Brown for animal husbandry.

## Notes

### Competing Interest Statement

The authors have declared no competing interest.

